# A widespread Xrn1-resistant RNA motif composed of two short hairpins

**DOI:** 10.1101/522318

**Authors:** Ivar W. Dilweg, Alexander P. Gultyaev, René C. Olsthoorn

## Abstract

Xrn1 is a major 5′-3′ exoribonuclease involved in the RNA metabolism of many eukaryotic species. RNA viruses have evolved ways to thwart Xrn1 in order to produce subgenomic non-coding RNA that affects the hosts RNA metabolism. The 3’ untranslated region of several beny-and cucumovirus RNAs harbors a so-called ‘coremin’ motif that is required for Xrn1 stalling. The structural features of this motif have not been studied in detail yet. Here, by using *in vitro* Xrn1 degradation assays, we tested over 50 different RNA constructs based on the Beet necrotic yellow vein virus sequence, to deduce putative structural features responsible for Xrn1-stalling. We demonstrated that the minimal benyvirus stalling site consists of two hairpins of 3 and 4 base pairs respectively. The 5’ proximal hairpin requires a YGAD (Y = U/C, D = G/A/U) consensus loop sequence, whereas the 3′ proximal hairpin loop sequence is variable. The sequence of the 9-nucleotide spacer that separates the hairpins is highly conserved and potentially involved in tertiary interactions. Similar coremin motifs were identified in plant virus isolates from other families including *Betaflexiviridae, Virgaviridae* and *Secoviridae* (order of the *Picornavirales*). We conclude that Xrn-stalling motifs are more widespread among RNA viruses than previously realized.

## Introduction

In order to counteract and cope with infection by RNA viruses, eukaryotic cells have evolved methods to process and degrade viral RNA. For instance, double-stranded RNA, which is formed during replication of positive-strand RNA viruses, can be processed through endolytic cleavage by ribonuclease III-family proteins into small interfering (si)RNA^1,2^. Such siRNAs are subsequently utilized in RNA-induced silencing complexes (RISC), followed by the cleavage of complementary viral RNA^3,4^. As a counterdefense, RNA viruses have evolved ways to interfere with RNA silencing. Viruses from diverse families encode so-called RNA silencing suppressors (RSSs) which can either sequester siRNAs, like p19 of tombusviruses^5^ or interact with protein components of RISC, like VP35 of Ebola virus^6^. While RSSs directly or indirectly prevent virus RNA breakdown, they may also be involved in fine-tuning host-virus interactions by regulating host transcriptional gene silencing (TGS) and post-transcriptional gene silencing (PTGS)^7,8^. Another way by which viruses can regulate host PTGS is demonstrated by flaviviruses like yellow fever virus, which employ structures in the 3’ untranslated region (UTR) of their RNA to stall the exoribonuclease Xrn1^9–11^. The latter process results in the production of Xrn1-resistant RNA (xrRNA) or small subgenomic flavivirus RNA (sfRNA) that may attenuate RNA silencing through interference with RNAi pathways^12,13^, interfere with translation^14^, and are required for achieving efficient pathogenicity^9,15,16^. On the other hand, Hepatitis-C virus and pestivirus RNAs have the ability to bind miRNAs, thereby interfering with Xrn1-mediated degradation and RNAi pathways as well^17,18^.

These xrRNAs are not exclusive to flaviviruses however. The plant-infecting diantho-, beny-and cucumoviruses produce a subgenomic RNA through the action of an Xrn1-like enzyme^14,19^. Furthermore, certain arenaviruses and phleboviruses harbor structures that can stall Xrn1 *in vitro*^20^. While elaborate tertiary structures are required to block Xrn1 progression in flavivirus and dianthovirus RNAs^19,21^, the role of RNA structure in the production of beny-and cucumovirus subgenomic RNAs has remained enigmatic. During infection of *Beta macrocarpa* by Beet necrotic yellow vein virus (BNYVV), a member of the *Benyviridae* family and *Benyvirus* genus^22^, a non-coding RNA is produced from BNYVV RNA3^23^. This RNA, and in particular the “core” sequence it carries, has been shown to be necessary for long-distance movement by the virus and can be produced by action of either yeast Xrn1 or plant XRN4^24–26^. A highly conserved 20 nucleotide (nt) sequence within the core, termed “coremin”, plays an important role in allowing for systematic infection by BNYVV RNA3 in *Beta macrocarpa*^23^. A recent study has indicated that these 20 nt are not sufficient to stall Xrn1 *in vitro*^26^ but that a minimum of 43 nt is required. Interestingly, the coremin motif is also found in the 3’ UTR of BNYVV RNA5, Beet soil-borne mosaic virus (BSBMV) and two species of cucumoviruses^23^. To date, it remains unknown whether RNA structure, like it does for xrRNAs in flaviviruses^21^, plays a role in this type of stalling.

In this study, we interrogate the coremin motif and flanking sequences for the requirement of secondary structure, thermodynamic stability and sequence conservation in achieving Xrn1 stalling. Over 50 RNA constructs were produced that systematically deviate in sequence throughout the expanded coremin motif. These constructs were subsequently tested for Xrn1 resistance *in vitro*. We show that Xrn1 resistance by the BNYVV RNA3 3’ UTR requires that the expanded coremin motif forms two stem-loop structures, one with a conserved and one with a variable loop sequence, which are separated from each other by a conserved spacer.

## Materials and Methods

### Prediction of coremin motif structure

The secondary structure of coremin motifs with various mutations or from various species was predicted *in silico* through the use of MFOLD^27^.

### PCR

Oligonucleotide templates representing different BNYVV 3’ UTR mutants were purchased from SigmaAldrich and Eurogentec in desalted form. Forward primers bear a T7 promoter sequence at the 5’ end. The 3’ ends of both forward and reverse primers carried reverse complementary sequences. A list of oligonucleotides is available on request. PCR reactions were carried out in a 50 µl volume, containing 400 nM of each oligo, 200 µM dNTPs and 2 units DreamTaq polymerase on a BioRad cycler. PCR fidelity was checked by agarose gel electrophoresis and products were purified by ethanol/NaAc precipitation at room temperature and dissolved in 25 µl Milli-Q water.

### *In vitro* transcription

*In vitro* transcription reactions were carried out using T7 RiboMAX™ Large Scale RNA Production System (Promega) in 10 µl volumes, containing 5 µl PCR product (∼250 ng), 5 mM of each rNTP, 1 µl Enzyme mix, in 1x Transcription Optimized buffer (40 mM Tris-HCl, 6 mM MgCl_2_, 2 mM spermidine, 10 mM NaCl, pH7.9 @ 25 ° C). After incubation at 37 ° C for 30 mins, 1 unit RQ1 RNase-Free DNAse was added to the reaction and incubation proceeded at 37 °C for 20 mins. Reaction samples were checked on agarose gel in order to establish subsequent usage of equal amounts of RNA.

### *In vitro* Xrn1 degradation assay

Xrn1 digestion reactions were performed with 1-4 µl RNA (∼400 ng, according to *in vitro* transcription yield) in 1x NEB3 buffer (100 mM NaCl, 50 mM Tris-HCl, 10 mM MgCl_2_, 1 mM DTT, pH 7.9 @ 25°C), totaling 10 µl, which was divided over two tubes. To one of the tubes, 0.2 units of Xrn1 and 0.3 units of RppH (New England Biolabs) were added. Both tubes were incubated for 15 mins at 37 ° C and the reactions were terminated by adding 5 µl formamide containing trace amounts of bromophenol blue and xylene cyanol FF. Samples were run on 14% native polyacrylamide gels in TAE buffer at 4 ° C using a MiniproteanIII system (BioRad) set at 140 V. Gels were stained with EtBr and imaged using a BioRad Geldoc system. Each construct was subjected to this assay at least twice.

## Results

### Phylogeny of the coremin motif

The alignment of coremin-containing 3’ UTR sequences from several viral species (Fig. 1) shows that the motif hairpin is very well conserved, as determined by others before^28^. Moreover, the BNYVV and CMV species harbor the coremin motif in multiple RNA species. Previously, the coremin motif was predicted to fold into a small hairpin^29^. At the 5’ and 3’ side of the motif, sequences are much more variable. Despite this, a recent study has shown that *in vitro* transcripts require at minimum 24 nt of the sequence 3’ of the BNYVV RNA3 coremin motif in order to safeguard Xrn1 resistance^26^. Structural analyses of the regions directly flanking coremin motifs in the aligned viral species using MFOLD^27^ identified no conserved structures 5’ of coremin but did reveal a putative hairpin structure 3’ of it. In most species, this hairpin (denoted here as hp2) is located directly after the conserved coremin motif hairpin (hp1). Between species, hp2 shows variable stem lengths and-composition, while the loops differ in size and sequence as well. However, structural alignment of hp2 (Fig. 1) reveals several instances of natural nucleotide covariation, which suggests a certain functionality for such coremin-flanking structures.

**Figure 1.**
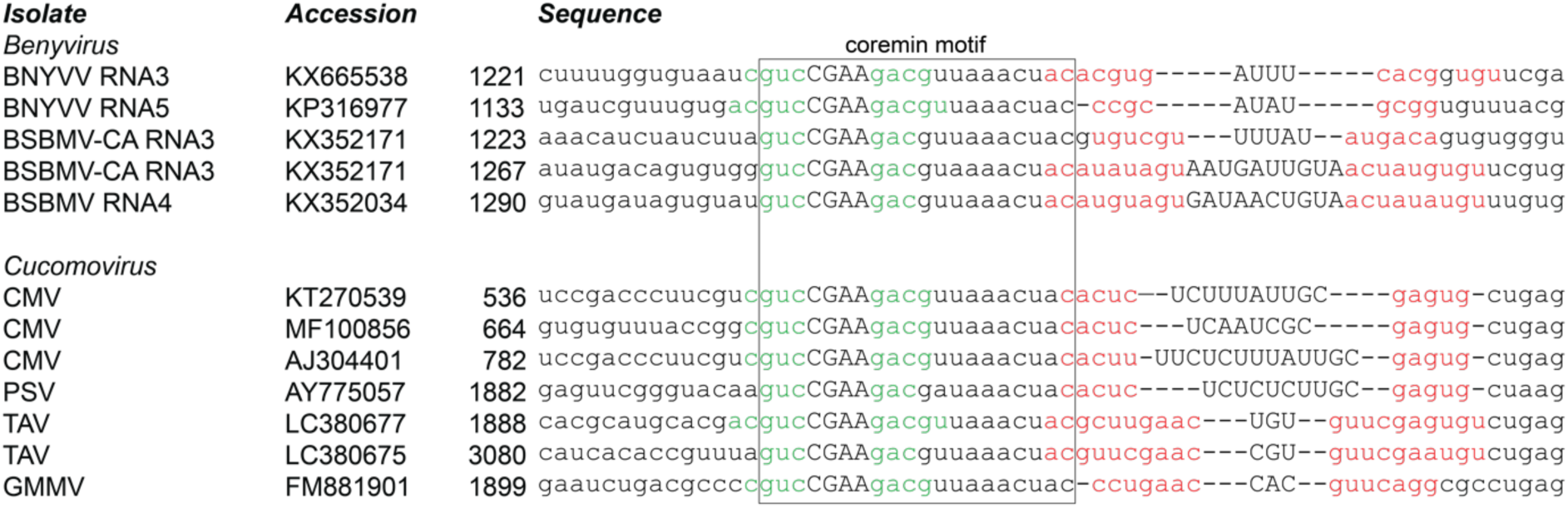
Alignment of coremin motifs in beny-and cucumoviruses. Multiple beny-and cucomovirus species harbor a coremin motif (boxed), which carries nucleotides that form a putative 3-5 bp-sized hairpin structure (green; uppercase letters depict the predicted loop sequence). An additional putative hairpin, more variable than, and directly downstream of the coremin motif, has been predicted by MFOLD for each sequence (base pairs formed by red nucleotides). Through structural alignment, covariations are revealed in this region. The position of the 5’ most nt is indicated for each sequence. RNA3 of BSBMV-CA isolate harbors two proximate coremin motifs. Note that this list is not exhaustive but shows the variation within these two genera. BSBMV: Beet soil-borne mosaic virus, CMV: Cucumber mosaic virus, PSV: Peanut stunt virus, TAV: Tomato aspermy virus, GMMV: Gayfeather mild mottle virus.

### Minimal construct for *in vitro* Xrn1 assays and role of hairpin 2

Based on the above findings we synthesized an RNA that comprises nucleotides 1224-1273 of BNYVV RNA-3 (NC°003516.1) preceded by a GA sequence for efficient transcription by T7 RNA polymerase. This RNA, when incubated with RppH (to generate the necessary 5’ monophosphate for Xrn1) and Xrn1, was processed to an RNA that had lost approximately 10 nt, showing that this construct is capable of efficiently stalling Xrn1 (Fig. 2, compare lanes “wt” plus and minus Xrn1). Truncating the RNA by 9 nt at its 3’ end (downs1) abolishes its stalling capacity, demonstrating that hp1 is not sufficient even though the coremin motif is not affected. A construct truncated by 5 nt (downs2) instead remained functional. Next, we tested whether nucleotide changes upstream of hp1 would influence Xrn1-resistance. To this end the GGUG sequence at positions 4-7 nt upstream of hp1 was changed to AAUA (ups). This change did not influence Xrn1 stalling and since the G-rich sequence could lead to unwanted alternative structures, AAUA variants were used for the majority of constructs in this study.

**Figure 2.**
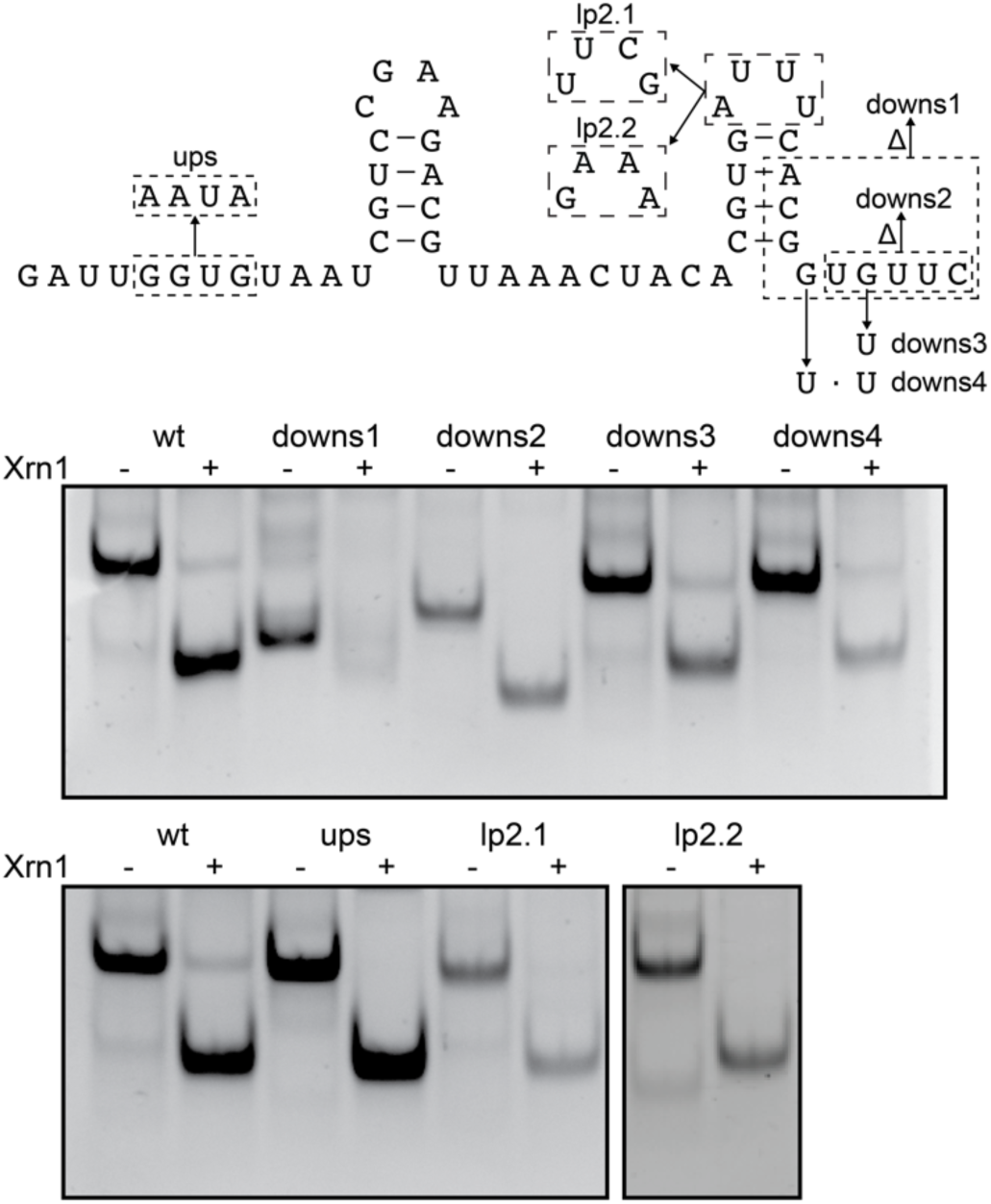
*In vitro* Xrn1 degradation assay probing requirement of sequence downstream of the coremin motif. Mutants used in this assay are imposed upon the BNYVV RNA3 construct depicted on top. RNA incubated with or without Xrn1 is loaded on native PAGE gels.

By replacing in the 3’ end of the wt construct either G35 (downs3) or both G35 and G37 (downs4) with U, we tested whether the stem-length of hp2 influences Xrn1 resistance. Both mutants remained able to stall Xrn1 in part, indicating that hp2 may consist of 4 or 5 bp, instead of 7 bp should A20C21A22 pair with U36G37U38 in the wild type construct. Moreover, replacing hp2 with a stable 9-bp hairpin still resulted in a construct that resisted complete degradation by Xrn1 (Supplementary Fig. 1). These mutants also show that nucleotides downstream of hp2 are not required and therefore, presumably not involved in interactions mediating structure. Due to the high variability observed in the hp2 loop (lp2) sequence of beny-and cucumoviruses, we expected that replacement of the loop by stable tetraloops would not affect resistance against degradation by Xrn1. Indeed, mutant constructs lp2.1 with UUCG^30^ and lp2.2 with GAAA^31^ were as resistant as wild type (Fig. 2).

### Role of spacer nucleotides

In order to assess whether nucleotides in the sequence linking hp1 and hp2 are crucial for Xrn1 resistance, several substitution mutants were designed (Fig. 3). Substituting U13U14 with AA (sp.sub1) did not affect Xrn1 stalling. In contrast, substitutions of A15A16A17 with UUU (sp.sub2), C18U19 with AA (sp.sub3) or UC (Supplementary Fig. 2) and A20C21 with UA (sp.sub4) all abolished Xrn1 stalling. The effects of these mutations were scrutinized more specifically through the investigation of their constituent single mutations. A17U (sp.sub6), C18A (sp.sub7), U19C (sp.sub8), U19A (sp.sub9) and A20U (sp.sub10) all resulted in constructs unable to resist Xrn1 as well. In contrast, A16U (sp.sub5), C21U (sp.sub11) and C21A (sp.sub12) mutations resulted in constructs that were roughly two-fold less resistant to Xrn1 than wt. These results indicate that the linker sequence, and in particular A17 until A20, fulfills an essential role within the coremin motif.

**Figure 3.**
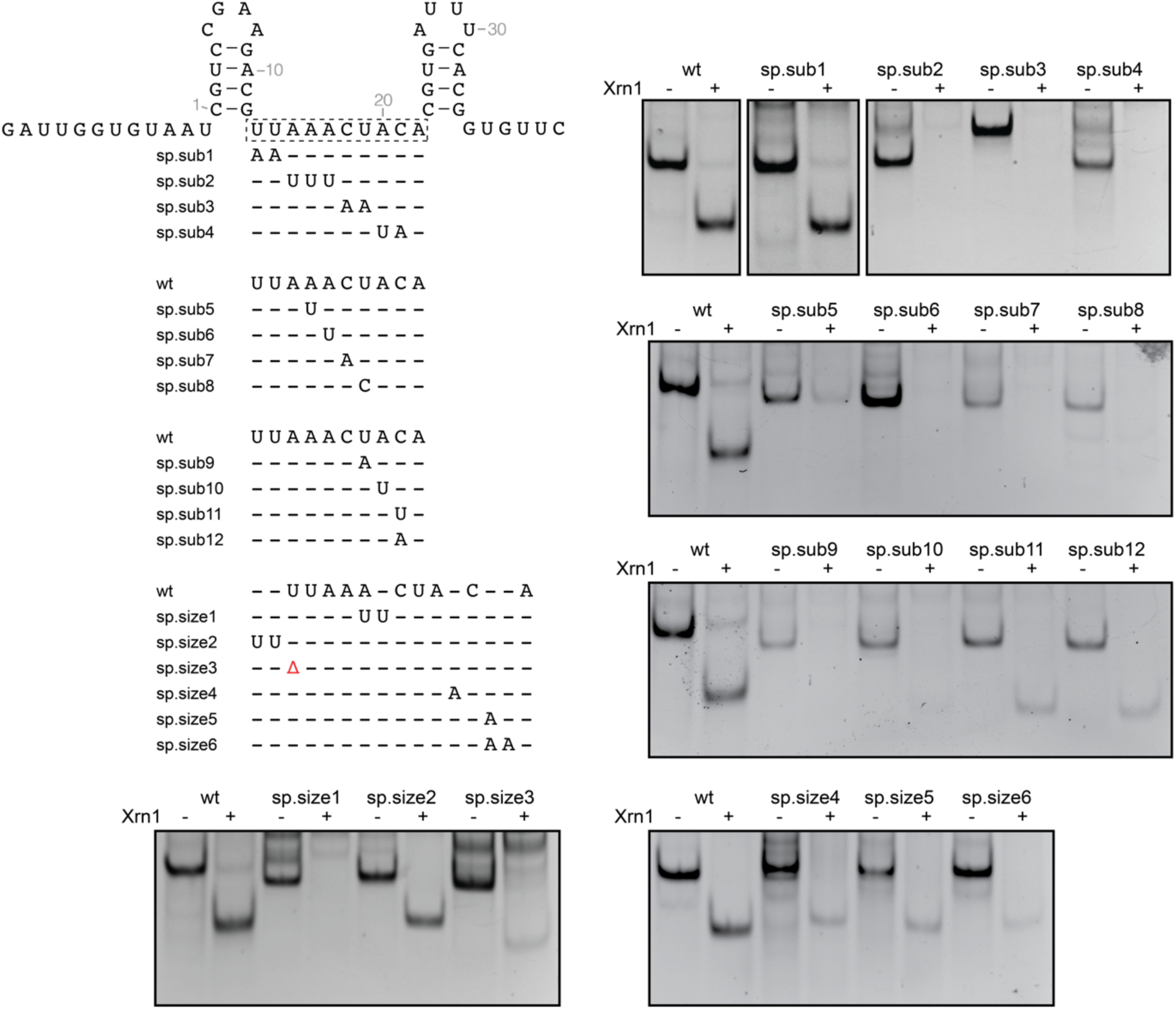
*In vitro* Xrn1 degradation assay aimed at the BNYVV RNA3 spacer sequence. Mutant constructs are depicted in alignment under the wildtype BNYVV RNA3 construct spacer sequence depicted on top. RNA incubated with or without Xrn1 is loaded on native PAGE gels.

In order to test whether the length of the spacer affects Xrn1 resistance, RNA was constructed carrying an insertion of UU after G12 (sp.size2). This only slightly reduced stalling of Xrn1. In contrast, a construct carrying an A17UU mutation (sp.size1) was not able to stall Xrn1 at all. Insertion of a single A after A20 (sp.size4) was not tolerated very well, as only a small fraction of RNA remained undegraded. Inserting either one (sp.size5) or two (sp.size6) adenosine residues after C21 also yielded such intermediate effects. Since the 3’ end of the spacer is apparently more sensitive towards mutations, insertions 3’ of either A20 or C21 may have disturbed potential interactions that these nucleotides undergo. A shorter spacer was tested as well, through deletion of U13 (sp.size3). This resulted in RNA that was degraded almost completely by Xrn1.

### Mutational analysis of hairpin 1

In contrast to hp2, the hairpin that forms the 5’ end of the conserved coremin motif (hp1) and its loop (lp1) shows much less variation in nature (Fig. 1). Previous experiments by Peltier *et al*. ^23^ demonstrated that changing lp1 to GACA is detrimental to Xrn1 resistance. We designed additional lp1 variations aimed at elucidating whether a certain structure or thermodynamic stability is required for Xrn1 stalling (Fig. 4). Out of thirteen lp1 mutants tested, only four were able to retain a level of Xrn1 resistance, namely UGAA (lp1.2), CGAU (lp1.8), CGAG (lp1.9) and, to a lesser extent, CAAA (lp1.3). These loops are not among those found to be very thermodynamically stable^32^, while conversely, the stable tetraloops GGAA^31^ (lp1.1), GAAA^31^ (lp1.11) and UUCG^30^ (lp1.12) do not yield Xrn1 resistant constructs. It is therefore likely that, in order to stall Xrn1, lp1 does not require a thermodynamically stable, but rather a certain conformation.

**Figure 4.**
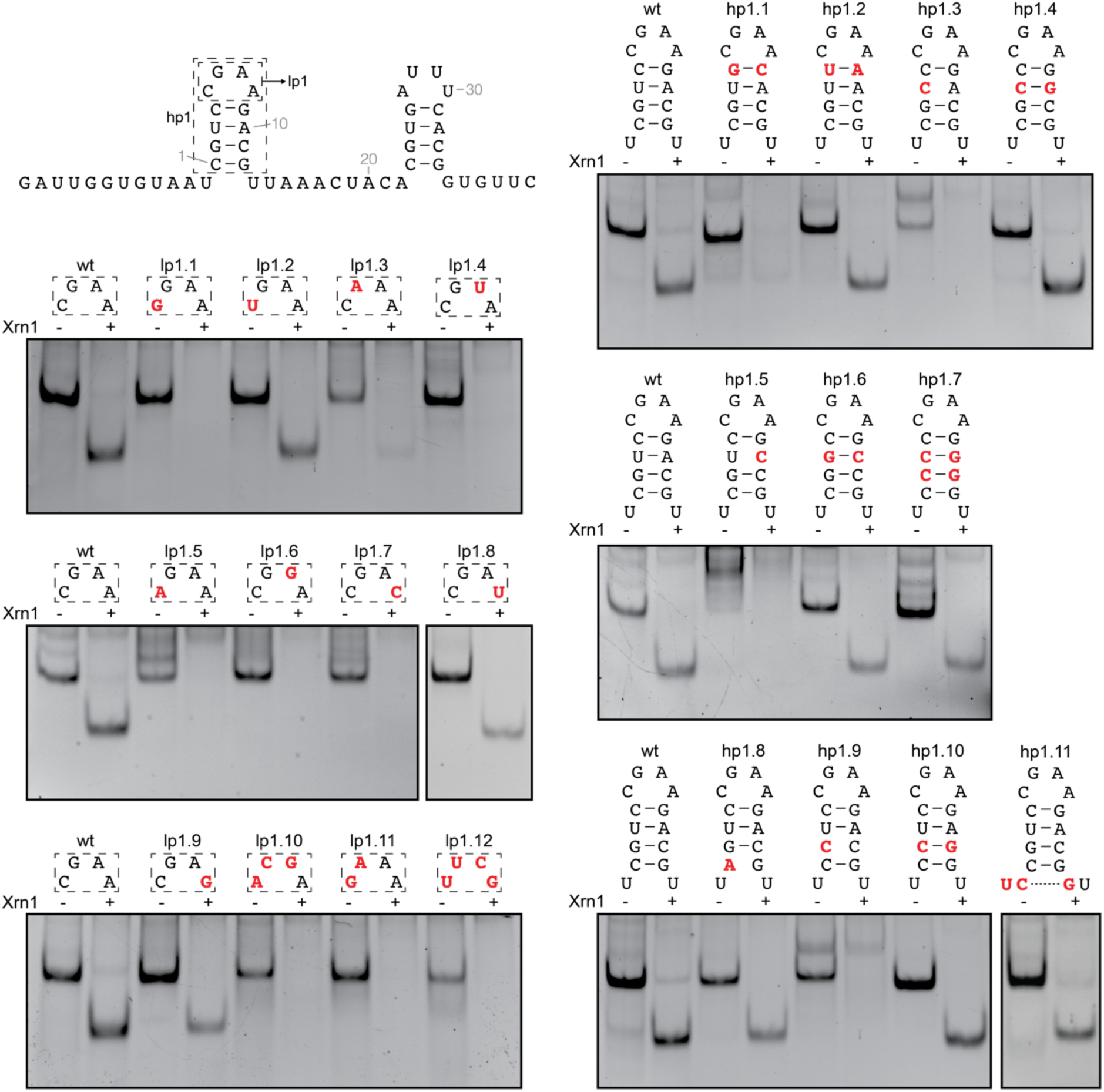
*In vitro* Xrn1 degradation assay targeted at the 5’ hairpin in BNYVV RNA3 coremin motif. Mutant constructs carrying either substitutions in the loop (lp1, left) or stem (hp1, right), are depicted above the corresponding native PAGE gels.

Additionally, we designed several mutants aimed to investigate the role of base pairing in the hp1 stem for Xrn1 stalling (Fig. 4). For the first base pair or loop closing base pair (lcbp) no disruption-restoration procedure was followed as lcbps are generally sequence specific and the above experiments showed that a certain loop conformation was required. Indeed, replacing it by a G-C bp (hp1.1) almost completely abolished Xrn1-resistance while a U-A bp (hp1.2) was slightly less resistant than wild type. Disruption of the second base pair by either a U3C (hp1.3) or A10C (hp1.5) mutation was found to abolish Xrn1-stalling. Restoring this base pair by a subsequent A10G (hp1.4) or U3G (hp1.6) mutation, respectively, also restored Xrn1-resistance. Similar effects were observed for the third base pair through disruption by G2 to C (hp1.9) and subsequent restoration by C11 to G (hp1.10). Moreover, a construct carrying C-G at each of the four base pairs (hp1.7), remained able to partially stall Xrn1.

Finally, through substituting C1 with A (hp1.8), the fourth base pair was disrupted, putatively resulting in a hairpin formed by three base pairs. This mutation only slightly reduced Xrn1 resistance. A potential fifth base pair can be formed by RNA5 of BNYVV as well as a sixth G-U bp. These extensions do not affect Xrn1 stalling as a transcript with the sequence of RNA5 as shown in Figure 1 remained as effective as our wild type (Supplementary Fig. 3). A hairpin of five base pairs in the context of BNYVV RNA3 has been tested through mutation of the AU directly on the 5’-side of the coremin to UC and U13 to G (hp1.11), which resulted in a construct able to stall Xrn1 as well. Together, these mutants indicate that within the stem of hp1, secondary structure is more important than sequence identity.

### Coremin-like sequences in other viral families

Although the coremin motif has been identified as a very well-conserved sequence, multiple nucleotide substitutions are tolerated by the motif, retaining the ability to stall Xrn1. Such variant sequences have been implemented for BLAST searches against ssRNA viruses in Genbank, which returned several novel hits. These putative xrRNAs were subjected to an *in vitro* Xrn1 degradation assay (Fig. 5).

**Figure 5.**
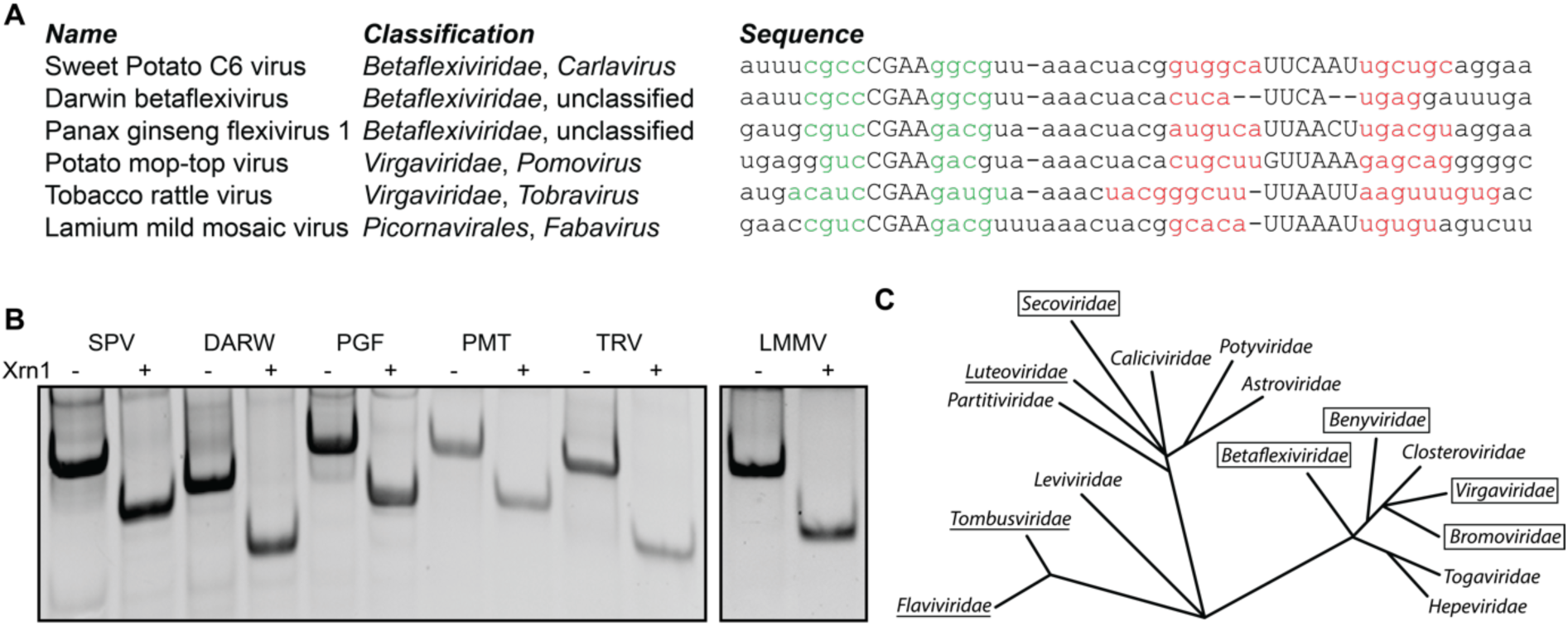
Widespread conservation of coremin motif in positive-strand ssRNA viruses. (A) List of novel coremin-like sequences found in different viral families. (B) *In vitro* Xrn1 degradation assay testing Xrn1-resistance for these sequences. (C) Phylogeny of positive-strand ssRNA viruses, based on RNA-dependent RNA polymerase conservation. Boxed viral families represent those containing viruses carrying a functional coremin-like sequence, including previously identified examples in *Bromoviridae*29. The *Flaviviridae, Tombusviridae* and *Luteoviridae* families are underlined, as they carry species with a non-coremin Xrn1 stalling site^10,14,33^. Phylogenetic tree is adapted from Koonin et al.^34^. SPV: Sweet Potato C6 virus, DARW: Darwin betaflexivirus, PGF: Panax ginseng flexivirus 1, PMT: Potato mop-top virus, TRV: Tobacco rattle virus, LMMV: Lamium mild mosaic virus.

1. A benyvirus isolate carrying CGAG in lp1 (KP316671). This corresponds with our mutant lp1.9, which turned out nearly as resistant as the wt CGAA lp1.
2. Two members of the *Betaflexiviridae* family, namely Sweet potato virus C-6 Sosa29 (JX212747) and Darwin betaflexivirus (MG995734), carrying a C-G as second bp in hp1, instead of the BNYVV RNA3 G-C. This natural covariation was tested using mutant hp1.10 and found capable of resisting Xrn1.
3. Another member of the *Betaflexiviridae* family, Panax ginseng flexivirus 1 (MH036372). This variant of coremin has a U14A substitution in the spacer. This sequence was found to be perfectly capable of stalling Xrn1.
4. An isolate of Potato mop-top virus (KU955493) from the Pomovirus genus within the *Virgaviridae* carrying a tandem repeat of the coremin motif in the 3’ UTR of its RNA-CP genomic segment. Such a repeated coremin motif has been identified in BSMBV-CA RNA3 as well (Fig. 1). Like Panax ginseng flexivirus 1, this variant of coremin carries a U14A substitution in the spacer.
5. Another member of the *Virgaviridae* family, Tobacco rattle virus (MF061245) from the Tobravirus genus, carrying a hp1 which deviates in both size (5 bp) and sequence from BNYVV RNA3 xrRNA. Moreover, this variant could putatively form a 9 bp hp2, incorporating more spacer nucleotides than seems to occur in other coremin-like xrRNAs.
6. The Lamium mild mosaic virus (KC595305), a Fabavirus belonging to the *Secoviridae* within the order of the *Picornavirales*, possessing one extra U in the spacer on the 3’ side of hp1. In context of BNYVV RNA3, we have shown that insertion of two uracils (sp.size2) at this position is tolerated as well (Fig. 3).

## Discussion

Previous studies on the 3’ UTR of both flavi-and dianthoviruses have indicated that elaborate structures are formed by the xrRNAs they utilize^19,21,35,36^. For instance, the crystal structure of Murray Valley Encephalitis Virus (MVE) flaviviral xrRNA revealed a ring-like conformation through tertiary interactions between its 5’ end and a downstream hairpin, which itself forms a pseudoknot with nucleotides even more downstream^36^. In doing so, a mechanical blockade is formed for Xrn1 that approaches the xrRNA from the 5’ end. Functional xrRNA derived from BNYVV RNA3 minimally requires fewer nt than that from the MVE flaviviral xrRNA. Therefore, there are fewer conformations possible that may result in stalling of Xrn1. We demonstrated here that xrRNA derived from the 3’ UTR of BNYVV RNA3 achieves Xrn1 resistance through two proximal hairpins, separated by a short spacer.

Although the RNA3 sequence forming hp1 is well-conserved, a few mutants targeted at this structure remained able to block Xrn1-mediated degradation. Changing the lcbp from C-G to U-A was tolerated, while switching the nucleotides to G-C abolished Xrn1-resistance almost entirely. This suggests that the specific loop conformation is favored by an upstream pyrimidine and downstream purine. Furthermore, substituting the U-A base pair, the second base pair from the top, with either G-C or C-G, or swapping the third base pair did not lead to loss of Xrn1-resistance. Moreover, through mismatching of the fourth base pair, we demonstrated that a hairpin formed by three base pairs stalls Xrn1 as well as wild type. Such a 3-bp hairpin is common in strains that harbor the coremin motif, as can be seen in Figure 1. In addition, since a 5-bp hp1 remains functional for Xrn1 stalling, as demonstrated by constructs hp1.11, sp.sub1 and RNA5, it can be concluded that the structural presence of hp1 is required, while its size and sequence identity are of lesser importance. This seems to contradict earlier findings on the accumulation of subgenomic CMV RNA5 by Thompson et al. (2008)^29^ who showed that disruption and subsequent restoration of hairpin base pairs all yielded a severe reduction in RNA5 levels after inoculation in plants. However, in their restored hairpin the lcbp became G-C, which does not stall Xrn1 and so resulted in complete degradation of the subgenomic RNA5 in their assays.

Several findings underline the need for the presence of the proposed second hairpin hp2, which has not been studied previously in benyviruses^23,26^, although a somewhat similar hairpin was proposed originally for subgenomic RNA accumulation of CMV RNA5^29^. Covariations found by alignment of several different species indicate that hp2 is likely structurally relevant, while its function in the context of Xrn1 resistance does not rely on its specific sequence. Each of the viral species carrying coremin motifs tested in Figure 5 carried substantially different second hairpins as well. Indeed, truncating the BNYVV RNA3 construct until C31, abolishing formation of hp2, renders it incapable of stalling Xrn1, while a shorter truncation indicates that this effect is not due to the loss of nucleotides downstream of the proposed hp2. Interestingly, the latter truncation, while in a different context, has been tested by Flobinus *et al*. ^26^ and was found to be unable to stall Xrn1, which led to their conclusion that more nucleotides of the RNA3 sequence are required at the 3’ end.

Most mutations targeted at the sequence linking hp1 and hp2 result in a complete loss of Xrn1-stalling capacity. Conservation of this linker sequence suggests that either some tertiary interaction may be required for Xrn1 resistance, or that either the sequence, or the structure that this sequence forms, is recognized by Xrn1 internally. As observed on the native PAGE gel in Figure 3, mutation of C18U19 to AA (sp.sub3) caused slower migration indicating conformational changes, which renders these nucleotides strong candidates for being involved in mediating some structural element. However, this should have become apparent from altered migration by either one of its constituent single mutants (sp.sub7 & sp.sub8). Current experimental conditions have not yielded such results. Nevertheless, mutations likely have a more destabilizing effect at the higher temperature during incubation with Xrn1, than at the lower temperature of native gel electrophoresis. Gel bands derived from control reactions lacking Xrn1 therefore may still retain their structure at native gel electrophoresis conditions. Alternatively, mutant constructs could remain structured until Xrn1 associates upstream and initiates its unwinding and degrading action^37^.

A pseudoknot-like interaction between the hp1 loop and spacer could confer the topology required for stalling Xrn1. The conserved nature of these regions, coupled with the fact that the 5’ end of the spacer tolerates insertions, while Xrn1 resistance is lost by a single nt deletion, are arguments that indeed point towards such a conformation. However, the exact interactions required for such a structure in our construct have not been identified yet. Switching C18 and U19 (sp.sub13), resulting in a sequence that could interact with G6 and A7 in a canonical anti-parallel fashion, did not yield Xrn1-resistant RNA. Furthermore, changing lp1 to CAGA in a mutant carrying UC instead of C18U19 (Supplementary Fig. 2) could not complement its loss of Xrn1-resistance. This result however, does not exclude the possibility of a pseudoknot-like interaction occurring, as other non-Watson-Crick interactions may be involved, and the CAGA lp1 may be topologically incompatible for this stringent formation. The role of the nucleotides linking hp1 and hp2 surpasses that of a spacer, as single, double or triple mutations across the sequence affect construct stability. The adenosine bases at positions 15-17 could not be mutated to uracils, combined nor individually, although their function remains unclear. In many tertiary structures, adenosine residues find their way in the minor groove of an adjacent helix, forming base triples with G-C base pairs, thus stabilizing this tertiary interaction^38–40^. Base triples play a crucial role in Xrn1-resistant RNAs of flaviviruses^21^ and dianthoviruses^19^. In addition to degradation assays, different approaches are necessary to elucidate the three-dimensional structure of coremin xrRNA.

We have demonstrated that novel coremin-like motifs can been found in the *Betaflexiviridae, Virgaviridae* and *Secoviridae* families. These results show that the coremin motif is more widespread among families of (plant) viruses than previously realized. Interestingly, members of the *Secoviridae* are closely related to *Luteoviridae*, a family in which recently novel dianthovirus-like xrRNAs have been discovered^33^. While the formation of these novel xrRNAs still has to be demonstrated *in vivo*, the contrast between such different types of xrRNA in apparently closely related species asks for investigation of their origin and function.

## References

1. Bernstein, E., Caudy, A. A., Hammond, S. M. & Hannon, G. J. Role for a bidentate ribonuclease in the initiation step of RNA interference. Nature 409, 363–366 (2001).

2. Lee, Y. et al. The nuclear RNase III Drosha initiates microRNA processing. Nature 425, 415–419 (2003).

3. Szittya, G., Molnár, A., Silhavy, D., Hornyik, C. & Burgyán, J. Short defective interfering RNAs of tombusviruses are not targeted but trigger post-transcriptional gene silencing against their helper virus. Plant Cell 14, 359–72 (2002).

4. Pantaleo, V., Szittya, G. & Burgyán, J. Molecular bases of viral RNA targeting by viral small interfering RNA-programmed RISC. J. Virol. 81, 3797–806 (2007).

5. Lakatos, L., Szittya, G., Silhavy, D. & Burgyán, J. Molecular mechanism of RNA silencing suppression mediated by p19 protein of tombusviruses. EMBO J. 23, 876–884 (2004).

6. Fabozzi, G., Nabel, C. S., Dolan, M. A. & Sullivan, N. J. Ebolavirus proteins suppress the effects of small interfering RNA by direct interaction with the mammalian RNA interference pathway. J. Virol. 85, 2512–23 (2011).

7. Zhao, J. H., Hua, C. L., Fang, Y. Y. & Guo, H. S. The dual edge of RNA silencing suppressors in the virus-host interactions. Curr. Opin. Virol. 17, 39–44 (2016).

8. Csorba, T., Kontra, L. & Burgyán, J. Viral silencing suppressors: Tools forged to fine-tune host-pathogen coexistence. Virology 479–480, 85–103 (2015).

9. Pijlman, G. P. et al. A Highly Structured, Nuclease-Resistant, Noncoding RNA Produced by Flaviviruses Is Required for Pathogenicity. Cell Host Microbe 4, 579–591 (2008).

10. Moon, S. L. et al. A noncoding RNA produced by arthropod-borne flaviviruses inhibits the cellular exoribonuclease XRN1 and alters host mRNA stability. Rna 18, 2029–2040 (2012).

11. Silva, P. A. G. C., Pereira, C. F., Dalebout, T. J., Spaan, W. J. M. & Bredenbeek, P. J. An RNA Pseudoknot Is Required for Production of Yellow Fever Virus Subgenomic RNA by the Host Nuclease XRN1. J. Virol. 84, 11395–11406 (2010).

12. Schnettler, E. et al. Noncoding flavivirus RNA displays RNA interference suppressor activity in insect and Mammalian cells. J. Virol. 86, 13486–500 (2012).

13. Moon, S. L. et al. Flavivirus sfRNA suppresses antiviral RNA interference in cultured cells and mosquitoes and directly interacts with the RNAi machinery. Virology 485, 322–329 (2015).

14. Iwakawa, H. et al. A Viral Noncoding RNA Generated by cis-Element-Mediated Protection against 5’->3’ RNA Decay Represses both Cap-Independent and Cap-Dependent Translation. J. Virol. 82, 10162–10174 (2008).

15. Manokaran, G. et al. Dengue subgenomic RNA binds TRIM25 to inhibit interferon expression for epidemiological fitness. Science (80-.). 350, 217–221 (2015).

16. Göertz, G. P. et al. Noncoding Subgenomic Flavivirus RNA Is Processed by the Mosquito RNA Interference Machinery and Determines West Nile Virus Transmission by Culex pipiens Mosquitoes. J. Virol. 90, 10145–10159(2016).

17. Conrad, K. D. et al. microRNA-122 Dependent Binding of Ago2 Protein to Hepatitis C Virus RNA Is Associated with Enhanced RNA Stability and Translation Stimulation. PLoS One 8, 1–11 (2013).

18. Mortimer, S. A. & Doudna, J. A. Unconventional miR-122 binding stabilizes the HCV genome by forming a trimolecular RNA structure. Nucleic Acids Res. 41, 4230–4240 (2013).

19. Steckelberg, A.-L. et al. A folded viral noncoding RNA blocks host cell exoribonucleases through a conformationally dynamic RNA structure. Proc. Natl. Acad. Sci. U. S. A. 201802429 (2018). doi:10.1073/pnas.1802429115

20. Charley, P. A., Wilusz, C. J. & Wilusz, J. Identification of phlebovirus and arenavirus RNA sequences that stall and repress the exoribonuclease XRN1. J. Biol. Chem. 293, 285–295 (2018).

21. MacFadden, A. et al. Mechanism and structural diversity of exoribonuclease-resistant RNA structures in flaviviral RNAs. Nat. Commun. 9, 1–11 (2018).

22. Gilmer, D., Ratti, C. & Consortium, I. R. ICTV Virus Taxonomy Profile: Benyviridae. J. Gen. Virol. 98, 1571–1572 (2017).

23. Peltier, C. et al. Beet necrotic yellow vein virus subgenomic RNA3 is a cleavage product leading to stable non-coding RNA required for long-distance movement. J. Gen. Virol. 93, 1093–1102 (2012).

24. Lauber, E., Guilley, H., Tamada, T., Richards, K. E. & Jonard, G. Vascular movement of beet necrotic yellow vein virus in Beta macrocarpa is probably dependent on an RNA 3 sequence domain rather than a gene product. J. Gen. Virol. 79, 385–393 (1998).

25. Flobinus, A. et al. A viral noncoding RNA complements a weakened viral RNA silencing suppressor and promotes efficient systemic host infection. Viruses 8, (2016).

26. Flobinus, A. et al. Beet necrotic yellow vein virus noncoding rna production depends on a 5’→3’ Xrn exoribonuclease activity. Viruses 10, 1–20 (2018).

27. Zuker, M. Mfold web server for nucleic acid folding and hybridization prediction. Nucleic Acids Res. 31, 3406–3415 (2003).

28. Ratti, C. et al. Beet soil-borne mosaic virus RNA-3 is replicated and encapsidated in the presence of BNYVV RNA-1 and -2 and allows long distance movement in Beta macrocarpa. Virology 385, 392–399 (2009).

29. Thompson, J. R., Buratti, E., de Wispelaere, M. & Tepfer, M. Structural and functional characterization of the 5’ region of subgenomic RNA5 of cucumber mosaic virus. J. Gen. Virol. 89, 1729–1738 (2008).

30. Cheong, C., Varani, G. & Tinoco, I. Solution structure of an unusually stable RNA hairpin, 5GGAC(UUCG)GUCC. Nature 346, 680–682 (1990).

31. Heus, H. A. & Pardi, A. Structural features that give rise to the unusual stability of RNA hairpins containing GNRA loops. Science 253, 191–4 (1991).

32. Bottaro, S. & Lindorff-Larsen, K. Mapping the Universe of RNA Tetraloop Folds. Biophys. J. 113, 257–267 (2017).

33. Steckelberg, A.-L., Vicens, Q. & Kieft, J. S. Exoribonuclease-Resistant RNAs Exist within both Coding and Noncoding Subgenomic RNAs. MBio 9, e02461–18 (2018).

34. Koonin, E. V. & Dolja, V. V. Virus World as an Evolutionary Network of Viruses and Capsidless Selfish Elements. Microbiol. Mol. Biol. Rev. 78, 278–303 (2014).

35. Chapman, E. G., Moon, S. L., Wilusz, J. & Kieft, J. S. RNA structures that resist degradation by Xrn1 produce a pathogenic dengue virus RNA. Elife 2014, 1–25 (2014).

36. Chapman, E. G. et al. The Structural Basis of Pathogenic Subgenomic Flavivirus RNA (sfRNA) Production. Science (80-.). 344, 307–310 (2014).

37. Jinek, M., Coyle, S. M. & Doudna, J. A. Coupled 5’ Nucleotide Recognition and Processivity in Xrn1-Mediated mRNA Decay. Mol. Cell 41, 600–608 (2011).

38. Jenkins, J. L., Krucinska, J., McCarty, R. M., Bandarian, V. & Wedekind, J. E. Comparison of a PreQ1riboswitch aptamer in metabolite-bound and free states with implications for gene regulation. J. Biol. Chem. 286, 24626–24637 (2011).

39. Nissen, P., Ippolito, J. A., Ban, N., Moore, P. B. & Steitz, T. A. RNA tertiary interactions in the large ribosomal subunit: The A-minor motif. Proc. Natl. Acad. Sci. 98, 4899–4903 (2001).

40. Pyle, A. M., Boudvillain, M. & de Lencastre, A. A tertiary interaction that links active-site domains to the 5’ splice site of a group II intron. Nature 406, 315–318 (2000).

