## Supplementary figures for "A widespread Xrn1-resistant RNA motif composed of two short hairpins"

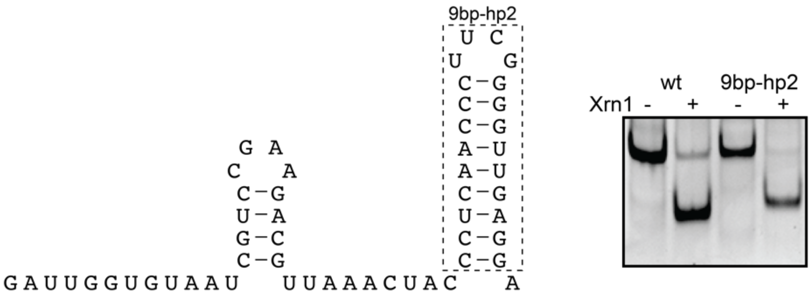


**Supplementary Figure S1.** ***In vitro*** **Xrn1 degradation assay shows that a 9-bp stem-loop can substitute for the wt BNYVV RNA3 coremin motif hp2.** The sequence on the left illustrates the putative secondary structure for the construct used in this experiment.

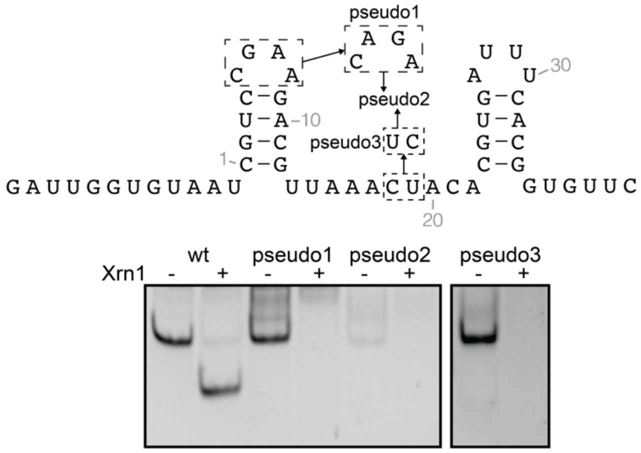


**Supplementary figure S2.** ***In vitro*** **Xrn1 degradation assay using mutants aimed at identifying a pseudoknot-like interaction between lp1 and the spacer.** Individually, substitution of lp1 with CAGA (pseudo1) and C18U19 with UC (pseudo3) result in loss of Xrn1 stalling capacity by the construct. Combining these mutations (pseudo2) did not recover Xrn1-resistance.

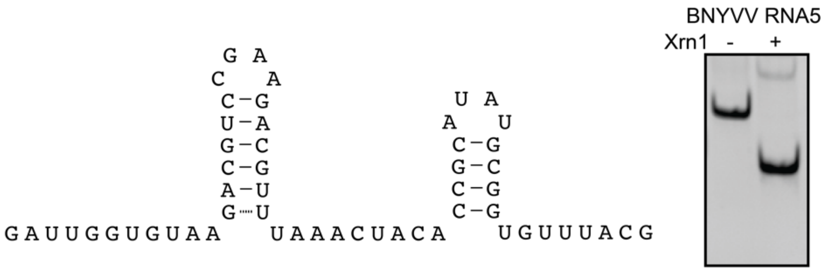


**Supplementary Figure S3. In vitro Xrn1 degradation assay demonstrates that BNYVV RNA5 carries a sequence that resists digestion by Xrn1.** The sequence on the left illustrates the putative secondary structure formed by the BNYVV RNA5 coremin motif.
